# Delayed dosing of minocycline plus N-acetylcysteine reduces neurodegeneration in distal brain regions and restores spatial memory after experimental traumatic brain injury

**DOI:** 10.1101/2021.03.28.437090

**Authors:** Kristen Whitney, Elena Nikulina, Syed N. Rahman, Alisia Alexis, Peter J. Bergold

## Abstract

Multiple drugs to treat traumatic brain injury (TBI) have failed clinical trials. Most drugs lose efficacy as the time interval increases between injury and treatment onset. Insufficient therapeutic time window is a major reason underlying failure in clinical trials. Few drugs have been developed with therapeutic time windows sufficiently long enough to treat TBI because little is known about which brain functions can be targeted if therapy is delayed hours to days after injury. We identified multiple injury parameters that are improved by first initiating treatment with the drug combination minocycline (MINO) plus N-acetylcysteine (NAC) at 72 hours after injury (MN72) in a mouse closed head injury (CHI) experimental TBI model. CHI produces spatial memory deficits resulting in impaired performance on Barnes maze, hippocampal neuronal loss, and bilateral damage to hippocampal neurons, dendrites, spines and synapses. MN72 treatment restores Barnes maze acquisition and retention, protects against hippocampal neuronal loss, limits damage to dendrites, spines and synapses, and accelerates recovery of microtubule associated protein 2 (MAP2) expression, a key protein in maintaining proper dendritic architecture and synapse density. These data show that in addition to the structural integrity of the dendritic arbor, spine and synapse density can be successfully targeted with drugs first dosed days after injury. Retention of substantial drug efficacy even when first dosed 72 hours after injury makes MINO plus NAC a promising candidate to treat clinical TBI.

## Introduction

Traumatic brain injury (TBI) is a major cause of death and disability world-wide (Faul and Coronado, 2015). Despite this critical need, decades of research have not produced FDA-approved pharmacological treatments for TBI (Diaz-Arrastia et al., 2014). TBI is an acute biomechanical event that produces a primary injury immediately upon impact and initiates a progressive secondary injury that develops over time (Blennow et al., 2012). The position of the hippocampus within the medial temporal lobe makes it particularly vulnerable to biomechanical injury (Bigler, 2013). The risk of long-term deficits increases as neural tissue is progressively damaged beyond the initial impact site (Bigler, 2013; Stocchetti and Zanier, 2016). As a result, TBI is best treated as early as possible. The majority of TBI’s, 70-80% of total incidence, are diagnosed as mild (Leibson et al., 2011). Despite the importance of early treatment, patients with a mild or moderate TBI often delay seeking treatment due to perceived symptom resolution, accessibility, or cost (Mohamadpour et al., 2019).

Most preclinical studies dose drugs within hours after injury, which does not consider the importance of treatment gap, the time between injury onset and treatment initiation (Tanielian, 2008; Demakis and Rimland, 2010). Treatment gap suggests that most TBIs will need to be treated with drugs with sufficiently long therapeutic time windows that target injury processes that develop over time (Diaz-Arrastia et al., 2014). Therapeutic time window in TBI is complex, as drug targets arise and disappear with differing kinetics (Mohamadpour et al., 2019). Despite the heterogeneity of TBI pathophysiology, most pharmaceutical interventions target a single injury mechanism (Xiong et al., 2013). Additionally, most preclinical studies have focused on treating injury in the hemisphere ipsilateral to the impact site. This focus may arise through the common use of unilateral open head experimental TBI models that directly injure the brain via a craniotomy (Failla and Wagner, 2015; Osier and Dixon, 2016). Neuronal loss and inflammation in open head models such as the controlled cortical impact model skew damage to regions proximal to the injury site. As a result, injury in regions distal to the impact site have been understudied (Hall et al., 2005; Tran et al., 2006). The closed head injury (CHI) model used in this study directly strikes an intact, unrestricted mouse skull with a piston. CHI produces a rapid acceleration-deceleration head movement resulting in a diffuse injury in both hemispheres (Johnson et al., 2015; White et al., 2017).

This study utilizes the diffuse brain injury produced by CHI to determine if the drug combination minocycline (MINO) plus N-acetylcysteine (NAC) retains efficacy when first dosed three days post-injury, and whether the therapeutic time window of MINO plus NAC differs in brain regions proximal or distal to the injury site. Both MINO and NAC have shown promise individually in preclinical experimental TBI models when first dosed within hours after injury (Xiong et al., 1999; Sanchez Mejia et al., 2001; Yi and Hazell, 2005; Hicdonmez et al., 2006; Bye et al., 2007; Chen et al., 2008; Abdel Baki et al., 2010; Homsi et al., 2010; Haber et al., 2017; Sangobowale et al., 2018). Although individual drugs alone showed reduced efficacy when first administered 24 hours after injury, MINO plus NAC synergized to lengthen the therapeutic time window in multiple outcomes when first dosed at 12 or 24 hours after CHI (Sangobowale et al., 2018). These data suggest that the drug combination of MINO plus NAC not only has higher efficacy, but also has a longer therapeutic time window than the individual drugs alone (Mohamadpour et al., 2019). MINO plus NAC restored acquisition of two hippocampal-dependent behavior tasks, active place avoidance and Barnes maze, and retained MAP2 and myelin levels in the ipsilateral hippocampus when dosed 12 hours after injury (Sangobowale et al., 2018). Unilateral hippocampal damage alone is not sufficient to induce spatial memory impairment. Lesion and inactivation studies demonstrate that spatial memory deficits need bilateral hippocampal destruction or inactivation (Smith et al., 1994; Poe et al., 2000; Tran et al., 2006). Injured mice first dosed 24 hours after injury (MN24) did not acquire active place avoidance, which requires two functional hippocampi, but did acquire Barnes maze, which only requires one functional hippocampus (Sunyer et al., 2007; Sangobowale et al., 2018). MN24 did not show histological improvement to the ipsilateral hippocampus, suggesting that the drug combination acted on the contralateral hippocampus to restore acquisition of Barnes maze.

Proper synapse formation and synaptic integration requires maintenance of the size, shape and complexity of dendritic arbors (Scott and Luo, 2001; Spruston, 2008). Dendrites, spines and synapses have high injury susceptibility since they are damaged by experimental TBI both ipsilateral and contralateral to the injury site (Scheff et al., 2005; Atkins, 2011; Gao and Chen, 2011; Gao et al., 2011; Campbell et al., 2012; Winston et al., 2013; Casella et al., 2014). Surviving neurons with damaged dendrites and fewer functional synapses results in disrupted brain circuitry. This study shows that damage to dendrites, spines and synapses is reduced by treatment with MINO plus NAC first dosed 3 days after CHI. Many of the therapeutic effects were seen in the contralateral hemisphere, suggesting regions distal to the initial injury can be targeted by drugs first dosed days after TBI.

## Methods

### Closed head injury model and drug treatment

All experiments were performed using male C57/BL6 mice (16 to 18 weeks old, 26-28gr). Mice were randomly assigned to control and experimental groups. Sham-CHI or CHI was administered as previously published (Grin’kina et al., 2016). Sham-injured mice received identical anesthesia treatment without the impact procedure, therefore the sham ipsilateral and contralateral hemispheres were combined. Mice were assessed for restoration of righting reflex before return to their home cage. Injured mice were randomly assigned to drug or saline treatment groups. At 72 hours after CHI or sham-CHI, mice received an intraperitoneal injection (i.p.) of MINO (22.5 mg/kg) plus NAC (75 mg/kg) in physiological saline or saline alone. Each group received two additional i.p. injections at 4 and 5 days post-injury. Sham-CHI mice only received saline treatment. All assessments done at 3 days after sham-CHI or CHI were on mice that did not receive drug or saline treatments. MINO plus NAC was prepared as previously described (Sangobowale et al., 2018). All drugs were from Sigma (St. Louis, MO). This study was carried out in strict accordance with the National Institutes of Health Guide for the Care and Use of Laboratory Animals. All studies were approved by the Institutional Animal Care and Use Committee of the State University of New York-Downstate Medical Center (protocol #18-10558). Experimenters were blinded to injury or treatment status since all mice were assigned a code number upon arrival at SUNY-Downstate that was used during all determinations.

### Barnes Maze

Behavioral assessment on Barnes maze (*n* =13-14/group) was performed as described by Sangobowale et al. (Sangobowale et al., 2018). Testing began seven days after sham-CHI or CHI, after transient motor deficits in injured mice return to sham-CHI levels (Grin’kina et al., 2016). For four consecutive days, each mouse received four 3-minute spatial acquisition trials with a 15-minute intertrial interval. On the 5^th^ day, mice received a 90-second probe trial with the escape box removed. Debut video software tracked the mouse position at 10 frames per second using a webcam (Tecknet C016 720p HD) located 1.5 m above the maze. Video records were analyzed using AnyMaze software (Stoelting) by an observer blind to the experimental treatment. The average of the four daily trials was calculated to produce the primary acquisition measure of latency to find the escape box. Latency, the number of erroneous holes searched, path length and speed were measured across the training trials. The probe trial was analyzed by time spent and number of holes searched in the target quadrant. Following behavioral assessment, animals were randomly assigned to histological analyses.

### Golgi-Cox staining and analysis

Fourteen days after sham-CHI or CHI, mice were anesthetized with urethane (0.1 mL 40%) and whole brains were stained as described in the FD Rapid Golgi Stain kit (FD Neurotechnologies, Columbia, MD, USA). Coronal sections (180μm) were prepared on a vibratome (Leica VT 100M), mounted and stained. Z-stacks of Golgi-stained dorsal hippocampus (from Bregma −1.3 to −2.7) were obtained with a Nikon DS-U3 camera on a Nikon Eclipse Ci-L microscope and processed with NIS-Elements D 4.40.00 software. Individual CA3 and CA1 pyramidal neurons were analyzed that had a fully impregnated dendritic tree that was minimally obscured by the dendrites of nearby neurons. CA1 and CA3 pyramidal cells meeting these criteria (10 cells per region) in each group (*n*=5 animals/group) were reconstructed into 3-dimensional traces using Neurolucida 360 software (Version 11.03, MBF Bioscience). NeuroExplorer software (MBF Bioscience) measured the reconstructed neurons for surface area and volume, branch morphology (total number, length, surface area and volume), and node number. Basal and apical dendritic trees were analyzed separately. Built-in convex hull analysis (NeuroExplorer, MBF Bioscience) measured neuronal volume and dendritic field size. Built-in Sholl analysis measured dendritic intersection number at 10μm intervals from the neuronal soma.

### Dendritic spine quantification and classification

Spines were quantified in CA1 apical dendrites proximal (<75 μm) and distal (>75μm) from the soma using NIS-Elements D 4.40.00 software. Secondary or tertiary dendritic branches were selected that did not overlap with other branches and were either parallel or at acute angles to the coronal section surface. Spine density per 10μm was assessed in at least 10 neurons and 50 branches per group (Table S3). Five distinct spine types (filopodia, stubby, thin, mushroom and branched) were manually sub-typed based on previously published methods (Arellano et al., 2007; Bloss et al., 2011; Risher et al., 2014; Afroz et al., 2016). Filopodia spines were defined as having a length of >2μm with no head; thin spines had a head/neck ration of >1:1 and a head diameter of <0.4μm; stubby spies had a length to width ratio of 1:1; mushroom spines had a head/neck ratio of >1:1 with a ≥0.4μm head width; and branched spine types contained 2 or more heads.

### Immunohistochemistry

Mice (*n*=3-5/group) were deeply anesthetized with isoflurane (3–5%) in oxygen (0.8 L/min) and fixed transcardially with paraformaldehyde (4%, w/v). Brain were post-fixed overnight in paraformaldehyde (4%, w/v), embedded in paraffin, and parasagittal sections (10μm) prepared 0.8-2mm from the midline (HistoWiz, Brooklyn, NY). Sections were stained with antibodies against MAP2 (Abcam, catalog #ab32454; antibody registry ID: AB 776174; rabbit polyclonal 1:500), NeuN (Abcam, catalog #ab104225; antibody registry ID: 10711153; rabbit polyclonal 1:500), synaptophysin (LifeSpanBioSciences, catalog #LS-C203763; antibody registry ID: AB 2864290; rabbit polyclonal, 1:500), and PSD-95 (Abcam, catalog #ab2723; antibody registry ID: 303248; mouse monoclonal, 1:500) with the appropriate fluorescent secondary antibodies. Nuclei were stained with 4′,6-diamidino-2-phenylindole. The specificity of each antibody binding was tested by staining adjacent regions with only primary or secondary antibodies. Sections were imaged using an Olympus Fluoview FV100 laser scanning confocal microscope using a 60x oil immersion objective and Leica Application Suite 1.8.2. Image J software measured MAP2 immunoreactivity in CA3 and CA1. Changes in MAP2 protein expression were examined separately in apical and basal dendritic trees and normalized to sham-CHI levels.

### Analysis of neuronal and synaptic density

Hippocampal cell density was quantified in NeuN-stained sections using stereological methods in NIH ImageJ software (Wang et al., 2003; Sun et al., 2006). NeuN immunopositive cells (NeuN^+^) were manually counted within each region in three 10μm sections beginning 0.8mm from Bregma. NeuN^+^ cell number in each section was divided by the product of the area of counting frame and section thickness, averaged and expressed as number of neurons per mm^3^. Synaptic density was quantified by staining intensity and colocalization of fluorescently labeled synaptophysin and PSD-95 using the method of Chakroborty et al. (Chakroborty et al., 2019) performed with Metamorph software (Version 7.1, Molecular Devices, Sunnyvale, CA). A median filter was applied to background-subtracted red and green imaging channels to reduce noise, and images were set to inclusive threshold levels. Equally sized image boxes (100 μm x 100 μm) were placed over the CA1 stratum radiatum. Staining intensity was determined as the percentage of area containing puncta of synaptophysin or PSD-95, over a threshold intensity level. Colocalization was measured using the Measure Colocalization plugin and expressed as the percentage of overlap of red and green fluorescent puncta. For each image stack, staining intensity and colocalization was measured in 3 serial planes and the results were averaged.

### Statistical Analysis

Two-way repeated measures (RM) ANOVA analyzed group differences in Barnes maze latency, errors, speed and path length on the factors of group and trial for each day assessed followed by pairwise comparisons using Tukey’s post-hoc test. A two-level mixed-effects model analyzed Sholl analysis on the factors of distance from the soma and treatment, followed by Tukey’s post hoc test. All remaining analyses used one-way ANOVA followed by pairwise comparisons using Tukey’s post hoc test. Statistical analysis was performed using Graphpad Prism software (La Jolla, CA) with significance set at 0.05. P-values in the text are from post-hoc comparisons unless noted otherwise as from ANOVA. All values are reported as mean ⩲ standard error of the mean.

## Results

### MN72 restores spatial navigation and memory deficits in Barnes maze after CHI

Beginning 7 days post-injury (PI), all groups were tested on Barnes maze. Latency had a significant effect of treatment (F_2,6_ = 62.36, p < 0.0001) and day (F_3,9_ = 101.9, p < 0.0001) with a significant interaction of treatment and day (F_6.18_ = 3.439, p < 0.05). Latency to find the escape hole significantly decreased from training days 1 to 4 in Sham-CHI and CHI-MN72 mice, but not CHI-Saline mice (Figure 1A). On Day 1, latency did not differ among the groups (p > 0.8). On Day 4, latency did not differ between Sham-CHI and CHI-MN72 mice and the latency of both groups was shorter than CHI-Saline mice (p < 0.005). Analysis of errors in Barnes maze testing yielded similar results (Table S1). Neither path length nor speed differed among the groups suggesting similar motor performance, therefore impaired navigation is the likely cause of differences in latency and errors (Table S1).

**Figure 1.**
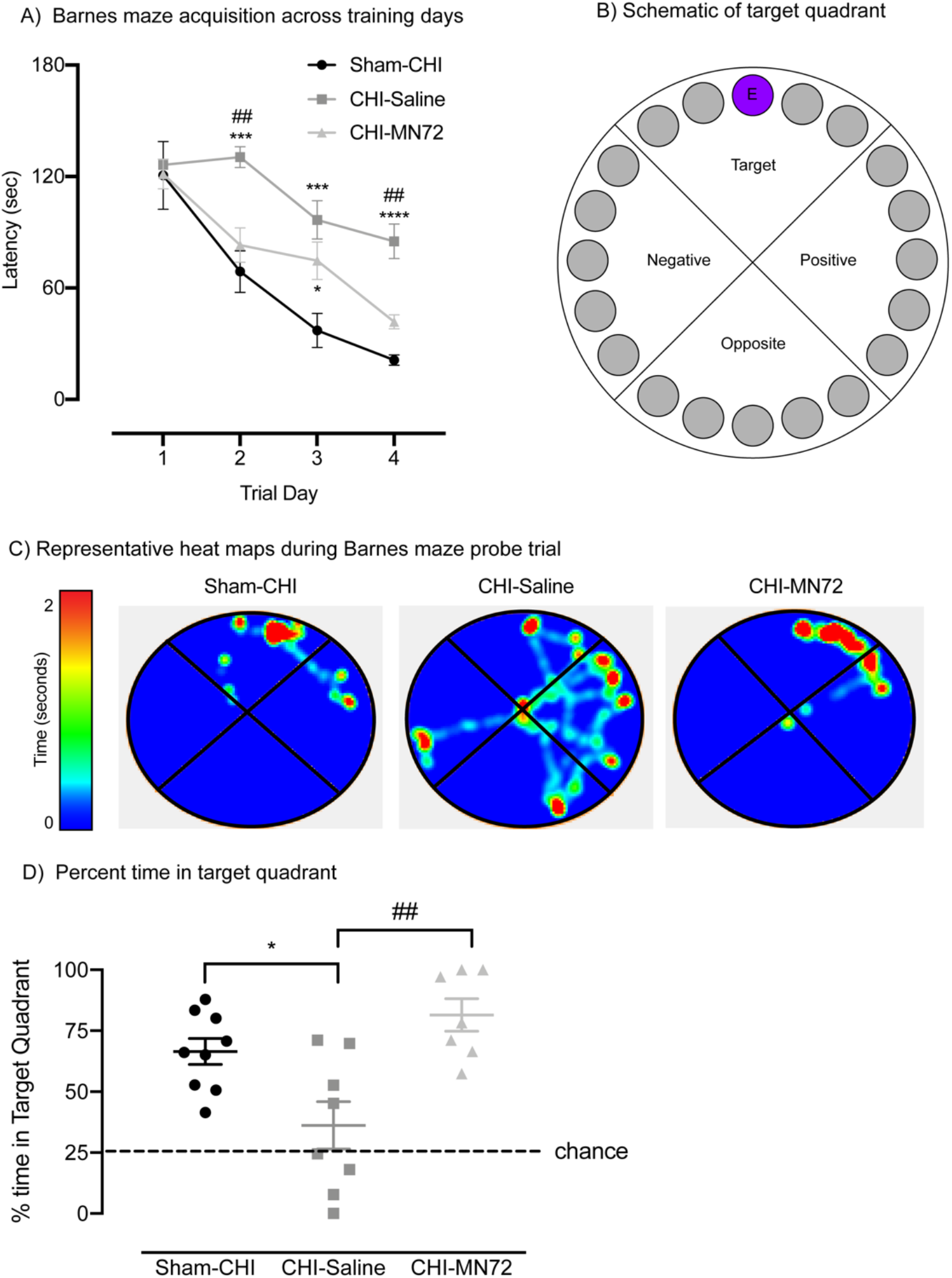
MN72 improves CHI-induced spatial navigation and memory deficits on Barnes Maze. **A**, CHI causes deficits on Barnes Maze acquisition. Group differences in latency are indicated across training days. On day 1, all groups had similar latencies. On day 4, Sham-CHI and CHI-MN72 mice had significantly shorter latencies than CHI-Saline mice. **B**, Schematic of Barnes maze with a target quadrant containing an escape hole (E). **C**, Representative heat maps of probe trials. **D**, Probe trial performance. Sham-CHI and CHI-MN72 mice spent more time in the target quadrant than CHI-Saline mice. A hash tag (#) indicates a significant difference with CHI-MN72 group, and an asterisk (*) indicates a significant difference with the Sham-CHI group; *p < 0.05, ***p<0.001, ****p<0.0001.

A probe trial assessed escape hole retention 24 hours after the last training trial (Figure 1B-D). The groups significantly differed in the percent time spent and holes searched in the target quadrant (% time, F_2,21_ = 9.35, p < 0.002; % holes, F_2,21_ = 13.24, p = 0.0002). Sham-CHI (p < 0.02) and CHI-MN72 mice (p < 0.002) spent significantly more time in the target quadrant than CHI-Saline mice (Figure 1D, Table S1). CHI-Saline mice searched significantly fewer holes than either Sham-CHI (p = 0.0002) or CHI-MN72 mice (p < 0.005; Table S1). Sham-CHI and CHI-MN72 mice did not differ in time spent or holes searched in the target quadrant (% time, p > 0.3; % holes, p > 0.7). These data strongly suggest that MN72 treatment restored the spatial navigation and memory deficits produced by CHI to uninjured levels.

### MN72 prevents neuronal loss in contralateral CA3 and ipsilateral CA1

Hippocampal neuronal loss after experimental TBI correlates to severity of hippocampal-dependent deficits on behavior tasks (Smith et al., 1997; Atkins, 2011). Therefore NeuN^+^ cells in CA1 and CA3 stratum pyramidale at 14 days PI were quantified using stereological methods (Figure 2A) (Wang et al., 2003; Sun et al., 2006). NeuN^+^ cell density in CA3 and CA1 had a significant group effect (CA3, F_4,17_ = 5.2, p < 0.007; CA1, F_4,15_ = 5.7, p < 0.006). CA3 NeuN^+^ density decreased bilaterally in CHI-Saline mice as compared to Sham-CHI (ipsilateral, p < 0.005; contralateral, p < 0.05; Figure 2B). In contrast, NeuN^+^ density decreased only in the ipsilateral CA3 of CHI-MN72 mice (ipsilateral, p < 0.05; contralateral, p > 0.2). CA1 NeuN^+^ density decreased in the ipsilateral, but not contralateral, hippocampus of CHI-Saline mice as compared to Sham-CHI mice (ipsilateral, p < 0.02; contralateral, p > 0.1; Figure 2C). In contrast, CA1 NeuN^+^ density did not differ in either hippocampus between Sham-CHI or CHI-MN72 mice (ipsilateral, p > 0.1; contralateral, p > 0.9). These data suggest that CHI produced CA3 neuronal loss in both hippocampi and CA1 neuronal loss in the ipsilateral hippocampus by 14 days PI. MN72 prevented CA3 neuronal loss in the contralateral hippocampus and CA1 neuronal loss in the ipsilateral hippocampus.

**Figure 2.**
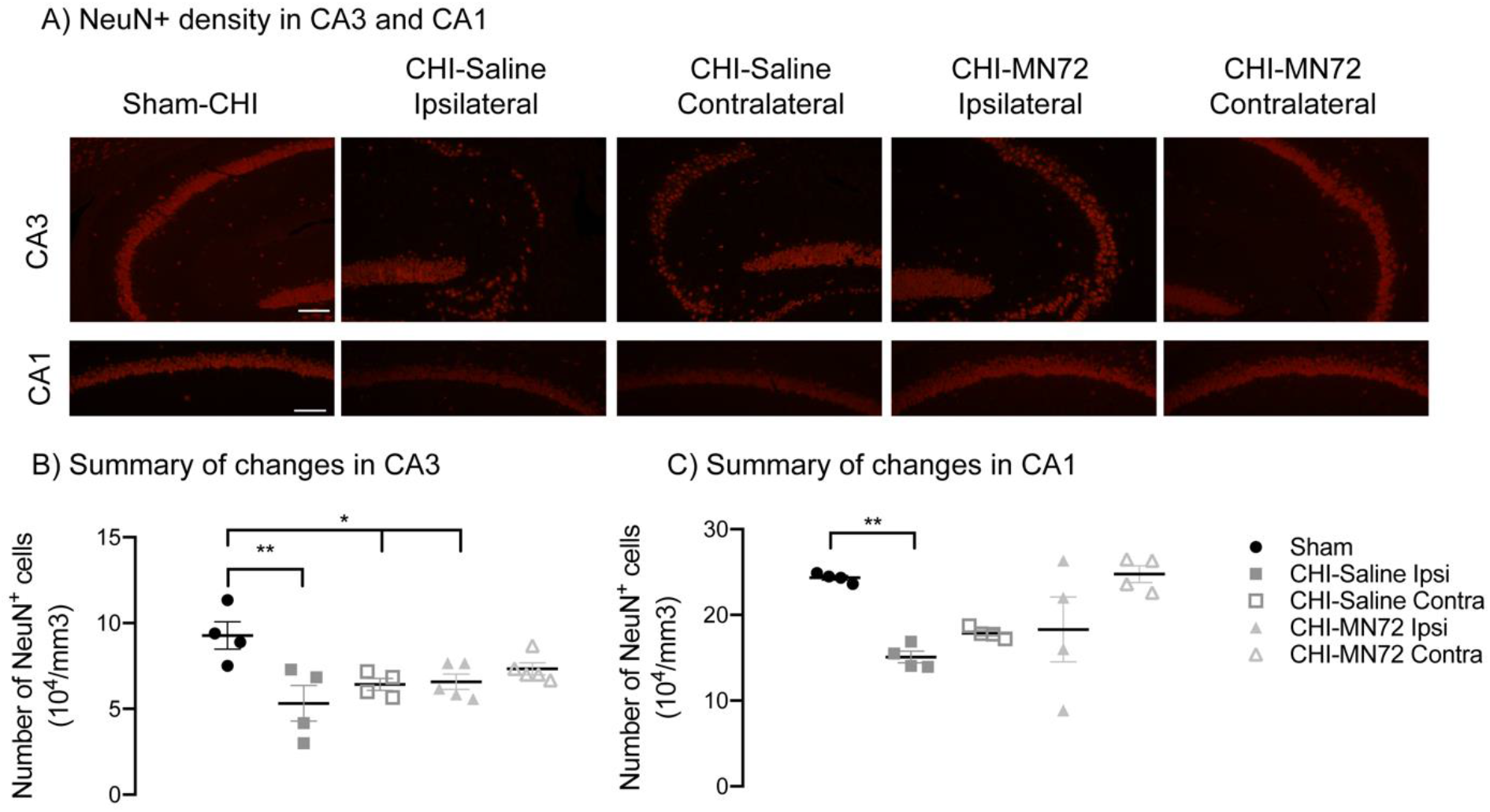
MN72 maintains neuronal density in contralateral CA3 and ipsilateral CA1. **A**, Representative photomicrographs of NeuN^+^ stained CA3 and CA1 at 14 days PI (scale bar, 100μm). **B**, Summary of CA3 NeuN^+^ density. Compared to Sham-CHI, CA3 NeuN^+^ density is bilaterally decreased in CHI-Saline mice, and decreased in the ipsilateral hippocampus in CHI-MN72 mice. **C,** Summary of CA1 NeuN^+^ density. Decreased ipsilateral CA1 NeuN^+^ density in CHI-Saline mice is prevented by MN72 treatment. An asterisk (*) represents significant differences with Sham-CHI, *p < 0.05, **p < 0.01.

### MN72 maintains CA3 and CA1 neuron volume and dendritic field size

Damage to neuronal and dendritic architecture produced by experimental TBI is more extensive than neuronal loss (Gao and Chen, 2011; Gao et al., 2011). Therefore, multiple morphological and geometric characteristics were assessed in Golgi-Cox stained CA1 and CA3 neurons in both hemispheres (Table S2). Convex hull analysis assessed the volume of surviving CA1 and CA3 neurons (Figure 3A, 3D). CA3 neuron volume differed among groups (F_4, 41_ = 8.09, p < 0.0001; Figure 3B). Bilaterally, the volume of CA3 neurons from CHI-Saline mice was reduced compared to Sham-CHI (ipsilateral, p < 0.005; contralateral, p < 0.01). In contrast, only CA3 neurons from the ipsilateral hippocampus of CHI-MN72 mice had reduced volume (p < 0.007). CA3 neuron volume in the contralateral hippocampus of CHI-MN72 mice did not differ from Sham-CHI (p > 0.5) and was larger than CHI-Saline mice (p < 0.05). These data suggest that CHI bilaterally reduced CA3 neuron volume, which was prevented in the contralateral hemisphere by MN72 treatment.

**Figure 3.**
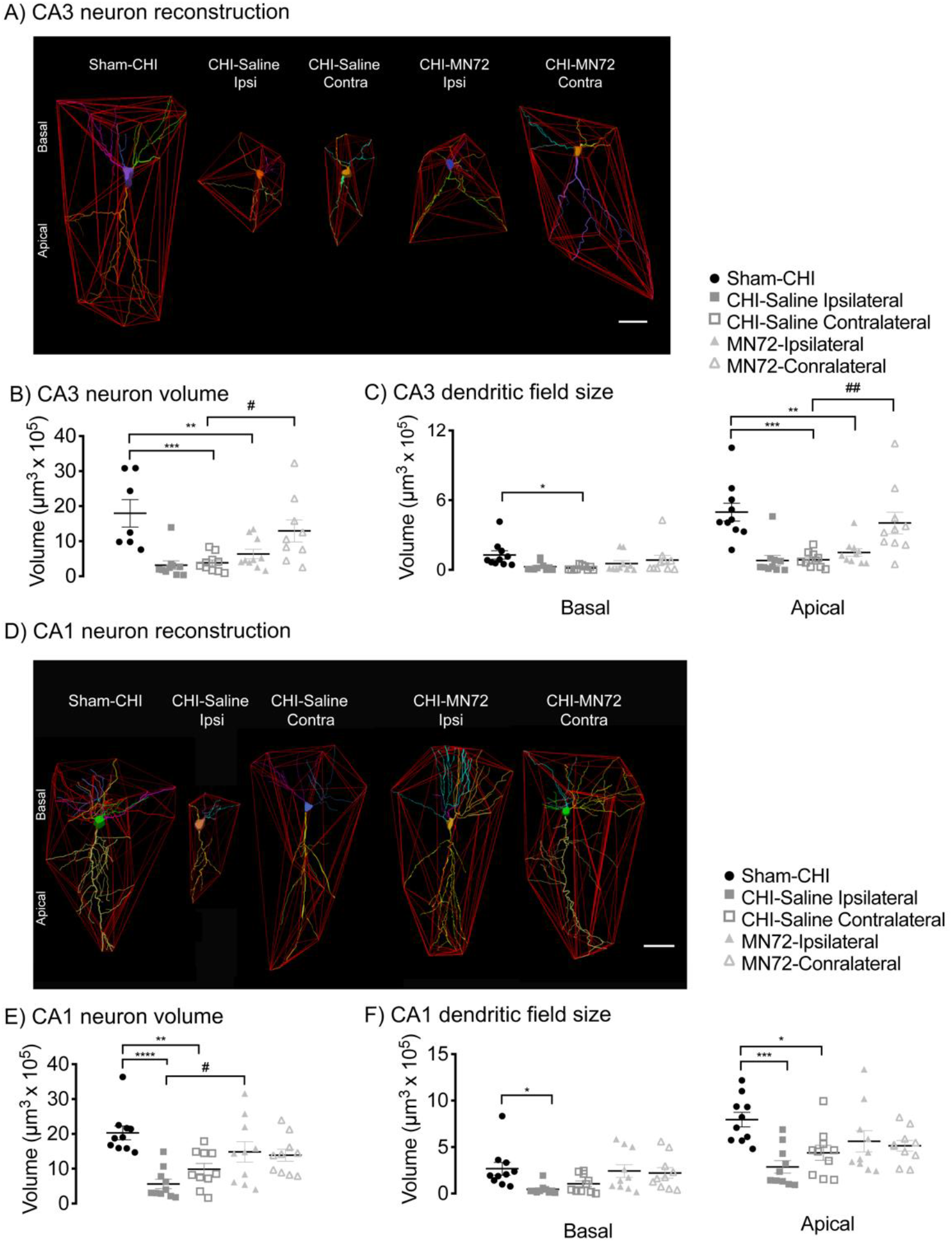
MN72 prevents reductions in CA3 and CA1 neuronal volume and dendritic field size induced by CHI. **A, D,** Polygons generated to assess CA3 (**A**) and CA1 (**D**) neuron volume and dendritic field size. **B**, CHI bilaterally reduced CA3 neuron volume which was prevented by MN72 in the contralateral hippocampus. **C,** CHI reduced contralateral CA3 basal dendritic field size and bilaterally reduced CA3 apical dendritic field size. MN72 prevented reduction of CA3 dendritic field size only in contralateral apical dendrites. **E**, CHI bilaterally reduced CA1 neuron volume which was prevented by MN72. **F**, MN72 bilaterally maintained CA1 basal and apical dendritic field sizes. An asterisk (*) represents significant difference with Sham-CHI, a hashtag (#) represents significant differences with CHI-MN72. * p < 0.05, ** p < 0.01, *** p < 0.001, **** p < 0.0001.

Convex hull analysis also measured basal and apical dendritic field size in CA3 and CA1 neurons. CA3 basal and apical dendritic field size significantly differed among groups (basal, F_4, 45_ = 2.79, p < 0.05; apical, F_4, 45_ = 10.32, p < 0.001; Figure 3C). In the ipsilateral hippocampus, CA3 basal dendritic field size was similar in CHI-Saline, CHI-MN72, and Sham-CHI mice. Ipsilateral apical dendritic field size was reduced in both CHI-Saline (p < 0.0002) and CHI-MN72 mice (p < 0.002). In the contralateral hippocampus, CA3 basal and apical dendritic field size in CHI-Saline mice was smaller than Sham-CHI mice (basal, p < 0.05; apical, p < 0.0005). In contrast, CA3 basal and apical dendritic field size was similar in CHI-MN72 and Sham-CHI mice (basal, p > 0.7; apical, p > 0.8). Apical dendritic field size was larger in CHI-MN72 and Sham-CHI mice than CHI-Saline mice (p < 0.005). These data suggest that, in CA3, CHI reduced basal dendritic field size in the contralateral hippocampus and reduced apical dendritic field size bilaterally. MN72 maintained CA3 apical dendritic field size in the contralateral hippocampus.

CA1 neuron volume also differed among groups (F_4, 45_ = 7.49, p = 0.0001; Figure 3E). Bilaterally, CA1 neurons from CHI-Saline mice had reduced volume compared to Sham-CHI mice (ipsilateral, p < 0.0001; contralateral, p < 0.005). In contrast, CA1 neurons from CHI-MN72 and Sham-CHI mice had similar volume (ipsilateral, p > 0.3; contralateral, p > 0.15). Ipsilateral CA1 neurons from CHI-MN72 mice had a larger volume than CHI-Saline mice (p < 0.02). These data suggest that MN72 bilaterally prevented the reduction of CA1 neuron volume produced by CHI.

CA1 basal and apical dendritic field size significantly differed among groups (basal, F_4, 45_ = 3.31, p <0.05; apical, F_4, 45_ = 5.17, p = 0.002; Figure 3F). In ipsilateral CA1, CHI-Saline mice had smaller basal and apical dendritic fields than Sham-CHI mice (basal, p < 0.05; apical, p < 0.01). In contralateral CA1, CHI-Saline mice had a similar basal but smaller apical dendritic field size (basal, p > 0.2; apical, p < 0.05). In contrast, CA1 basal and apical dendritic field size did not differ between CHI-MN72 or Sham-CHI mice in either hippocampus (basal, ipsilateral, p > 0.9, contralateral, p > 0.9; apical, ipsilateral, p > 0.2, contralateral, p > 0.1). These data suggest that CHI bilaterally reduced CA1 basal and apical dendritic field size, which was prevented by MN72.

### MN72 maintains CA3 dendritic architecture in the contralateral hippocampus

The shape and complexity of the dendritic arbor are important determinants of neuronal input and synaptic integration and could underlie functional deficits (Ishizuka et al., 1995; Staff et al., 2000; Stuart and Spruston, 2015). Dendritic architecture was therefore evaluated by two complementary methods, quantifying nodes and Sholl analysis (Figures 4,5; Table S2). Node number indicates the complexity of the dendritic arbor; Sholl’s analysis provides additional spatial information on dendritic arborization that enhances node analysis.

**Figure 4.**
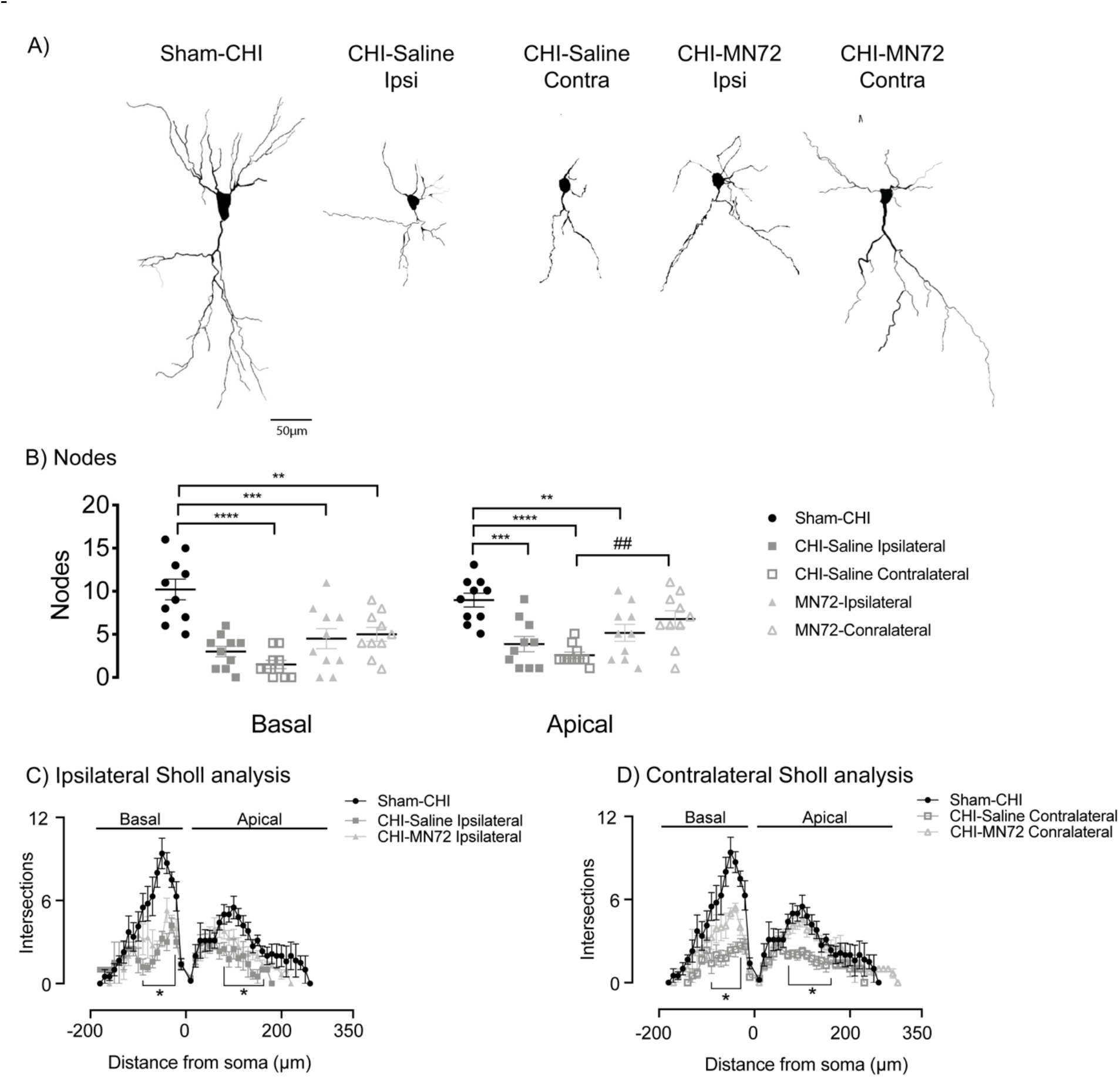
MN72 maintains CA3 dendritic complexity and architecture in the contralateral hippocampus. **A** Representative tracings of Golgi-Cox stained CA3 pyramidal neurons from Sham-CHI, CHI-Saline and CHI-MN72 mice. **B,** CHI bilaterally reduced node number in apical and basal dendrites compared to Sham-CHI. MN72 maintained node number in the contralateral CA3 apical dendrites. An asterisk (*) represents significant difference with Sham-CHI, a hashtag (#) represents significant differences with CHI-MN72. ** p < 0.01, *** p < 0.001, **** p < 0.0001. **C-D**, Sholl’s analysis. **C**, In the ipsilateral hippocampus, intersection number was reduced in CHI-Saline basal (30-90μm) and apical (80-160μm) dendrites compared to Sham-CHI. Intersection number was only lower in CHI-MN72 basal (30-60μm) dendrites. **D,** In the contralateral hippocampus, intersection number was reduced in CHI-Saline basal (30-70μm) and apical (70-160μm) dendrites compared to Sham-CHI. CHI-MN72 mice had fewer intersections in basal (30-50μm) dendrites than Sham-CHI mice, but more intersections (40-50μm) than CHI-Saline mice. Contralateral apical dendrites of Sham-CHI and CHI-MN72 mice had a similar number of intersections. An asterisk (*) indicates significant differences in intersection number.

**Figure 5.**
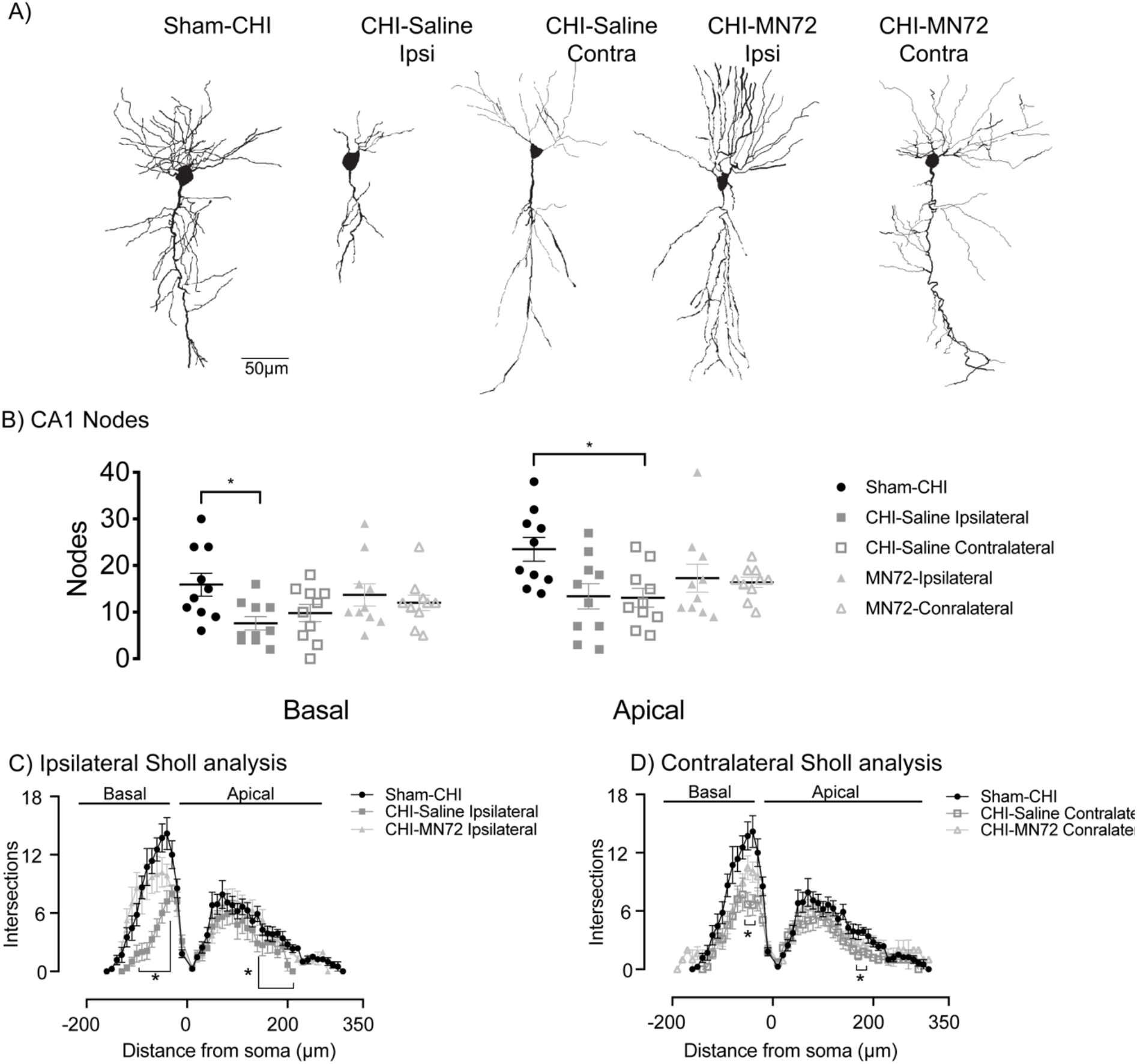
MN72 maintains CA1 dendritic complexity and architecture in the ipsilateral and contralateral hippocampus. **A** Representative tracings of Golgi-Cox stained CA1 pyramidal neurons from Sham-CHI, CHI-Saline and CHI-MN72 mice. **B,** CHI reduced node number in ipsilateral basal dendrites and in ipsilateral and contralateral apical dendrites. MN72 bilaterally maintained node number in both apical and basal dendrites. An asterisk (*) represents significant difference with Sham-CHI. * p < 0.05. **C-D**, Sholl’s analysis. **C,** In the ipsilateral hippocampus compared to Sham-CHI, intersection number was reduced in CHI-Saline basal (40-100μm) and apical (140-220μm) dendrites. Sham-CHI and CHI-MN72 mice had similar intersection number in ipsilateral basal and apical dendrites. **D,** In the contralateral hippocampus compared to Sham-CHI, intersection number was reduced in CHI-Saline basal (40-50μm) and apical (170-180μm) dendrites. Sham-CHI and CHI-MN72 mice had similar intersection number in contralateral basal and apical dendrites. An asterisk (*) indicates significant differences in intersection number.

Node number in CA3 basal and apical dendrites significantly differed among groups (basal, F_4, 45_ = 13.4, p < 0.0001; apical, F_4, 45_ = 9.01, p < 0.0001; Figure 4B). Bilaterally, apical and basal dendrites in CHI-Saline mice had fewer nodes than Sham-CHI mice (basal, ipsilateral, p < 0.0001, contralateral, p < 0.0001; apical, ipsilateral, p = 0.007, contralateral, p < 0.0001). In the ipsilateral hippocampus, apical and basal dendrites in CHI-MN72 mice had fewer nodes than Sham-CHI mice (basal, p < 0.002; apical, p < 0.05). In the contralateral hippocampus, CHI-MN72 mice had fewer nodes in basal, but not apical dendrites (basal, p < 0.002; apical, p > 0.3). Contralateral CA3 apical dendrites in CHI-MN72 mice had more nodes than CHI-Saline mice (p < 0.008).

Sholl’s analysis investigated dendritic architecture as a function of distance from the soma by assessing the number of dendritic crossing intersections with concentric circles radiating from the soma (Figures 4C,D). Treatment and distance had a significant group effect on intersection number (basal, treatment, F_4, 45_ = 12.79, p <0.0001, distance, F_3.2, 145.2_ = 58.17, p < 0.0001, treatment x distance, F_68, 765_ = 4.12, p < 0.0001; apical, treatment, F_4, 45_ = 14.52, p <0.0001, distance, F_3.9, 174.9_ = 51.66, p < 0.0001, treatment x distance, F_116, 1305_ = 2.79, p < 0.0001; Figures 4C, D). CHI bilaterally reduced intersection number across the CA3 basal and apical dendritic arbor compared to Sham-CHI (p < 0.05). MN72 treatment increased intersection number at discrete locations in the contralateral basal dendritic arbor (p <0.01; Figure 4D). Despite this improvement from CHI-Saline mice, CA3 basal dendrites in CHI-MN72 mice had fewer intersections than Sham-CHI mice near the soma in both hippocampi (p < 0.05). In contrast, in both ipsilateral and contralateral CA3 apical dendrites, intersection number did not differ between Sham-CHI and CHI-MN72 mice (p > 0.1). MN72 treatment also increased intersection number at discrete locations in the contralateral apical dendritic arbor compared to CHI-Saline mice (p < 0.05). These data suggest that CHI bilaterally altered branch distribution and reduced the complexity of CA3 apical and basal dendritic architecture. MN72 bilaterally maintained dendritic architecture with a greater therapeutic effect in contralateral CA3 dendrites.

### MN72 maintains CA1 dendritic architecture in the ipsilateral and contralateral hippocampus

Total node number in CA1 basal and apical dendrites significantly differed among groups (basal, F_4, 45_ = 2.64, p < 0.05; apical, F_4, 45_ = 3.14, p < 0.05; Figure 5B). In ipsilateral CA1 dendrites, CHI-Saline mice had fewer nodes compared to Sham-CHI (basal, p < 0.05; apical, p < 0.05). In contralateral CA1, CHI-Saline mice had fewer nodes in apical, but not basal dendrites (basal, p > 0.2; apical, p < 0.05). In contrast, node number in MN72-treated mice did not differ with Sham-CHI mice in either hippocampus (basal, ipsilateral, p > 0.9, contralateral, p > 0.6; apical, ipsilateral, p > 0.3, contralateral, p > 0.2).

In Sholl analysis, treatment and distance had a significant group effect on intersection number in CA1 dendrites (basal, treatment, F_4, 47_ = 6.89, p = 0.0002, distance, F_2.973, 139.7_ = 120.6, p < 0.0001, treatment x distance, F_72, 846_ = 3.93, p < 0.0001; apical, treatment, F_4, 48_ = 3.76, p = 0.01, distance, F_3.7, 171.6_ = 75.67, p < 0.0001; treatment x distance, F_120, 1380_ = 1.18, p < 0.09; Figure 5C,D). CHI bilaterally reduced intersections across CA1 apical and basal dendrites compared to Sham-CHI (p < 0.05). MN72 treatment increased intersection number across the CA1 apical and basal dendritic arbor compared to CHI-Saline mice (p < 0.05). In both hippocampi, intersection number did not differ between Sham-CHI and CHI-MN72 mice (basal, p > 0.08; apical, p > 0.1). These data suggest that CHI bilaterally altered CA1 dendritic branch distribution and reduced complexity with more pronounced injury in the ipsilateral hippocampus. MN72 bilaterally maintained CA1 dendritic architecture.

### MN72 either prevents MAP2 loss or accelerates MAP2 recovery in CA3 dendrites

MAP2 is a major dendritic cytoskeletal protein; loss of MAP2 expression accompanies dendritic damage in multiple TBI models (Taft et al., 1992; Posmantur et al., 1996; Grin’kina et al., 2016; Sangobowale et al., 2018). Lowered MAP2 expression induces dendritic instability and subsequent loss; both instability and loss can be restored by maintaining MAP2 expression. MN72 treatment preserves dendritic architecture (Figures 4, 5). We therefore examined a time course of changes in hippocampal MAP2 expression at 3, 7 and 14 days PI. CA3 pyramidal neurons in Sham-CHI mice had prominent MAP2 dendritic labeling and dendrites that projected orthogonally from CA3 stratum pyramidale into stratum lucidum and stratum radiatum (Figure 6A). In contrast, pyramidal neurons from CHI-Saline mice had diminished dendritic labeling and disorganized, non-linear dendrites. Contralateral CA3 pyramidal neurons from CHI-MN72 mice had restored dendritic labeling and linear organization.

**Figure 6.**
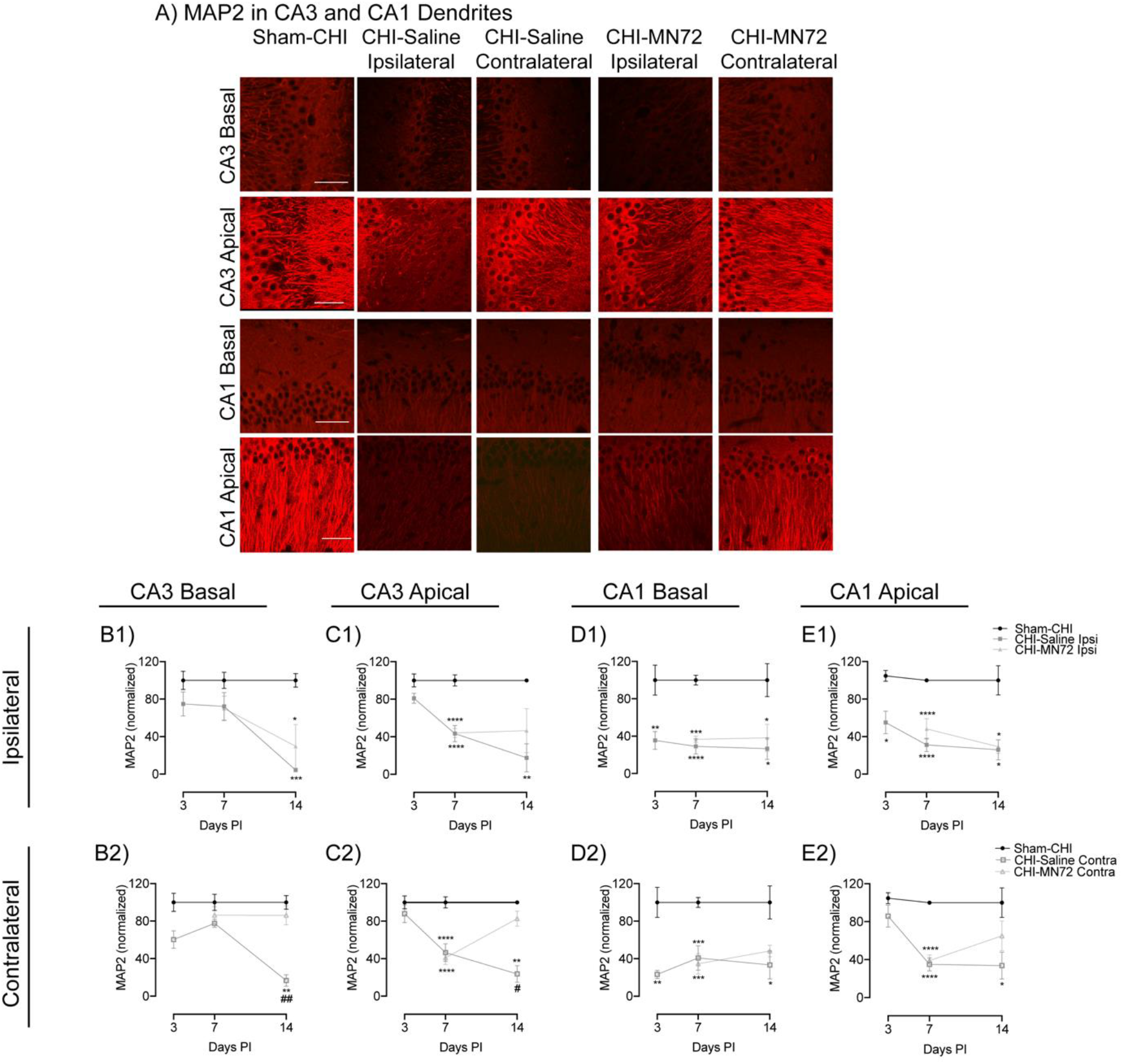
MN72 either prevents MAP2 loss or accelerates MAP2 recovery. **A**, Representative photomicrographs of MAP2 immunoreactivity in CA3 and CA1 apical and basal dendrites at 14 days PI. Scale bar, 50μm. Summary of changes in MAP2 expression in the ipsilateral (**B1, C1, D1, E1**) or contralateral (**B2, C2, D2, E2**) hippocampus. In all regions, MAP2 expression was unchanged in Sham-CHI mice. **B1,** In ipsilateral CA3 basal dendrites, MAP2 loss occurs between days 7-14 in CHI-Saline and CHI-MN72 mice. **B2,** In contralateral CA3 basal dendrites, MAP2 loss occurs between days 7-14 in CHI-Saline mice. MN72 treatment prevents MAP2 loss. **C1,** In ipsilateral CA3 apical dendrites, MAP2 loss occurs between days 3-7 in CHI-Saline and CHI-MN72 mice, and persists through days 7-14 in CHI-Saline mice. **C2**, In contralateral CA3 apical dendrites, MAP2 loss occurs between days 3-7 in CHI-Saline and CHI-MN72 mice, and persists through days 7-14 in CHI-Saline mice. In contrast, MAP2 expression increases between days 7-14 in CHI-MN72 mice. **D1**, In ipsilateral CA1 basal dendrites MAP2 loss occurs by day 3 and persists through day 14 in CHI-Saline and CHI-MN72 mice. **D2,** In contralateral CA1 basal dendrites, MAP2 loss occurs by day 3 in CHI-Saline mice and persists through day 14. In CHI-MN72 mice, MAP2 loss increases by day 14. **E1,** In ipsilateral CA1 apical dendrites, MAP2 loss by day 3 persists through day 14 in CHI-Saline and ChI-MN72 mice. **E2,** In contralateral CA1 apical dendrites, MAP2 loss occurs by day 7 in CHISaline and CHI-MN72 mice. MAP2 loss persists through day 14 in CHI-Saline mice. In contrast MAP2 is restored by day 14 in CHI-MN72 mice. An asterisk (*) represents a significant difference with Sham-CHI, a hashtag (#) represents significant difference with CHI-MN72. *p < 0.05, **p< 0.01. ***p < 0.001, ****p < 0.0001.

The time course of MAP2 expression in CA3 basal dendrites had a significant group effect only at 14 days PI (Day 3, F_2,8_ = 3.07, p > 0.1; Day 7, F_4,11_ = 1.52, p > 0.2; Day 14, F_4,11_ = 13.82, p = 0.003, Figure 6B1, B2). MAP2 expression in CHI-Saline and CHI-MN72 mice was unchanged at 3 and 7 days PI as compared to Sham-CHI (Day 3, CHI-Saline, ipsilateral, p > 0.3, contralateral, p > 0.08; Day 7, CHI-Saline, ipsilateral, p > 0.3, contralateral, p > 0.5; CHI-MN72, ipsilateral, p > 0.2, contralateral, p > 0.8). At 14 days PI, MAP2 decreased bilaterally in CHI-Saline mice, and ipsilaterally in CHI-MN72 mice (CHI-Saline, ipsilateral, p = 0.001; contralateral, p < 0.002; CHI-MN72, ipsilateral, p < 0.02; contralateral, p > 0.9). Contralateral MAP2 in CHI-MN72 mice was significantly higher than in CHI-Saline mice (p < 0.01). These data suggest that MN72 prevented MAP2 loss in CA3 basal dendrites in the contralateral hippocampus.

In CA3 apical dendrites, MAP2 expression had significant group effects at 7 and 14 days PI (Day 3, F_2,9_ = 1.68, p > 0.2; Day 7, F_4, 31_ = 12.96, p < 0.001; Day 14, F_4,11_ = 7.59, p < 0.005; Figure 6C1, C2). At 7 days PI, MAP2 decreased bilaterally in CHI-Saline and CHI-MN72 mice as compared to Sham-CHI (p < 0.0001). At 14 days PI, bilateral MAP2 loss persisted in CHI-Saline mice but not in CHI-MN72 mice (CHI-Saline, ipsilateral, p < 0.01, contralateral, p < 0.01; CHI-MN72, ipsilateral, p > 0.09; contralateral, p > 0.8). Contralateral MAP2 in CHI-MN72 mice was significantly higher than CHI-Saline mice (p < 0.05). These data suggest that MN72 induced MAP2 recovery in contralateral CA3 apical dendrites, while preventing further MAP2 loss in ipsilateral CA3 apical dendrites.

### MN72 accelerates MAP2 recovery in contralateral CA1 apical and basal dendrites

CA1 pyramidal neurons from Sham-CHI mice had prominent MAP2 labeling in long, linearly arranged dendrites (Figure 6A). In contrast, CHI-Saline mice had diminished MAP2 labeling in disorganized, non-linear dendrites. CHI-MN72 mice had increased MAP2 dendritic labeling and improved linear organization in the contralateral CA1 dendrites.

MAP2 expression in CA1 basal dendrites had significant group effects at 3, 7, and 14 days PI (Day 3, F_2,8_ = 15.66, p < 0.002; Day 7, F_4, 20_ = 13.52, p < 0.0001; Day 14, F_4,20_ = 4.090, p < 0.02, Figure 6D1, D2). At 3 days PI, MAP2 decreased bilaterally in CHI-Saline mice as compared to Sham-CHI (ipsilateral, p < 0.006, contralateral, p < 0.002). At 7 days PI, MAP2 decreased bilaterally in CHI-Saline and CHI-MN72 mice (CHI-Saline, ipsilateral, p < 0.0001, contralateral, p < 0.0004; CHI-MN72, ipsilateral, p < 0.0002, contralateral, p < 0.0003). MAP2 loss in CHI-Saline mice persisted bilaterally through 14 days PI (ipsilateral, p < 0.02, contralateral, p < 0.02). In contrast, MAP2 increased in the contralateral dendrites of CHI-MN72 mice (ipsilateral, p < 0.05, contralateral, p > 0.1).

In CA1 apical dendrites, MAP2 had significant group effects at 3, 7, and 14 days PI (Day 3, F_2,10_ = 5.79, p < 0.05; Day 7, F_4,31_ = 17.65, p < 0.0001; Day 14, F_4,19_ = 4.69, p < 0.009, Figure 6E1, E2). In Sham-CHI mice, MAP2 in CA1 apical dendrites was unchanged between days 3 and 14. At 3 days PI, ipsilateral MAP2 expression decreased in CHI-Saline mice as compared to Sham-CHI (ipsilateral, p < 0.02, contralateral, p > 0.4). At 7 days PI, MAP2 decreased bilaterally in both CHI-Saline and CHI-MN72 mice (p < 0.0001). Bilateral MAP2 loss in CHI-Saline mice persisted at 14 days PI (ipsilateral, p < 0.03, contralateral, p < 0.02). In contrast, at 14 days PI, CHI-MN72 mice had decreased MAP2 only in ipsilateral dendrites (ipsilateral, p < 0.05; contralateral, p > 0.4). These data suggest that MN72 accelerated MAP2 recovery in CA1 apical and basal dendrites in the contralateral hippocampus.

### MN72 maintains total and mushroom spine density in CA1 apical dendrites

Experimental TBI reduces spine density in CA1 apical dendrites with a severity that varies with the distance from the soma (Winston et al., 2016). Spines receive different presynaptic inputs depending upon their location from the CA1 pyramidal cell soma. In CA1 apical dendrites, Schaffer collateral axons synapse on proximal dendrites and perforant path axons synapse on distal dendrites (Spruston, 2008). Loss of Schaffer collateral or perforant path synapses could lead to Barnes maze deficits, therefore both proximal and distal CA1 apical dendrites were examined for spine loss following CHI (Figure 7A, B). Total spine density on proximal and distal dendrites differed significantly among groups (proximal, F_4,49_ = 7.73, p < 0.0001; distal, F_4, 50_ = 5.54, p < 0.001; Figure 7 C, Table S3). CHI bilaterally lowered total spine density on both proximal and distal dendrites (proximal, ipsilateral, p < 0.002, contralateral, p < 0.008; distal, ipsilateral, p < 0.02, contralateral, p < 0.005). Total proximal and distal spine density in Sham-CHI and CHI-MN72 mice did not differ in either hippocampus (proximal, p > 0.8; distal, p > 0.4). In the ipsilateral hippocampus, CHI-MN72 mice had a higher proximal spine density than CHI-Saline mice (p < 0.005). In the contralateral hippocampus, CHI-MN72 mice had a higher distal spine density (p < 0.02).

**Figure 7.**
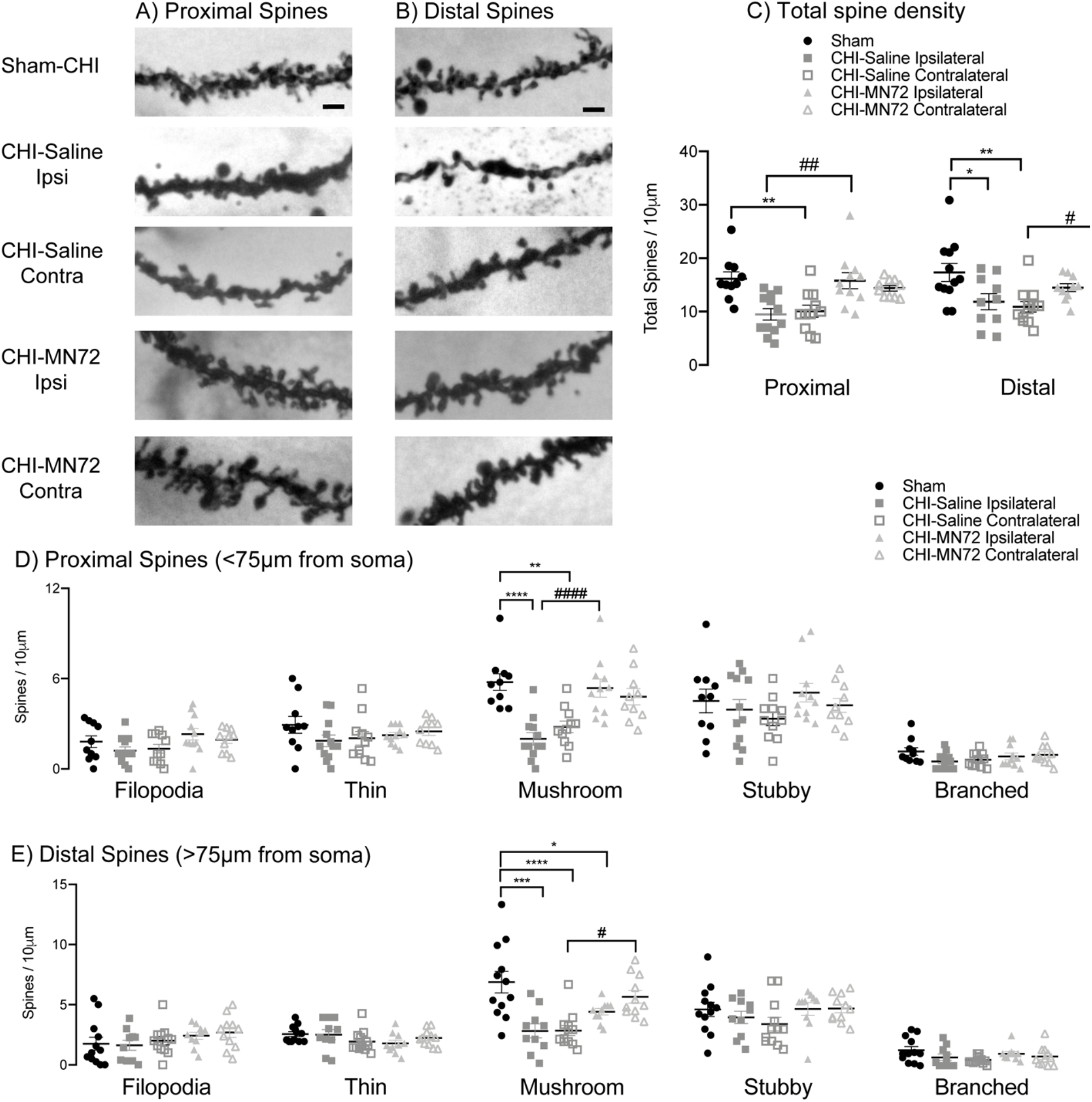
MN72 maintains spine density in CA1 apical dendrites. **A-B**, Representative images of proximal (<75 μm from soma) (**A**) and distal (>75 μm from soma) (**B**) Golgi-Cox stained CA1 apical dendrites. **C**, Summary of differences in total spine density. CHI bilaterally reduced total spine density in proximal and distal dendritic regions, which was maintained bilaterally by MN72. **D**, Individual spine type density in proximal dendrites. CHI bilaterally reduced proximal mushroom spine density. MN72 treatment maintained mushroom spine density. **E**, Individual spine type density in distal dendrites. CHI bilaterally reduced distal mushroom spine density. MN72 treatment maintained distal mushroom spine density in the contralateral hippocampus. An asterisk (*) represents significant a difference with Sham-CHI, a hashtag (#) represents a significant difference with CHI-MN72; *p < 0.05, ** p < 0.01, *** p < 0.001, ****p < 0.0001. Scale bar, 2μm.

Individual spine types were also assessed on proximal and distal dendrites. Mushroom spine density on proximal and distal dendrites differed among groups (proximal, F_4,49_ = 11.36, p <0.0001; distal, F_4,50_ = 9.05, p <0.0001; Figure 7D, E). CHI bilaterally decreased mushroom spine density on proximal and distal dendrites as compared to Sham-CHI (proximal, ipsilateral, p < 0.0001, contralateral, p < 0.002; distal, ipsilateral, p < 0.0005, contralateral, p < 0.0001). Bilaterally, mushroom spine density did not differ on proximal dendrites in Sham-CHI and CHI-MN72 mice (p > 0.6; Figure 7D). CHI-MN72 mice had more mushroom spines on proximal dendrites in the ipsilateral hippocampus than CHI-Saline mice (p < 0.001); increased mushroom spines in the contralateral hippocampus trended toward significance (p = 0.055). In contrast, Sham-CHI and CHI-MN72 mice had similar mushroom spine density on distal dendrites only in the contralateral hippocampus (ipsilateral, p < 0.05; contralateral, p > 0.5; Figure 7E). In the contralateral hippocampus, distal mushroom spine density was higher in CHI-MN72 mice than in CHI-Saline mice (p < 0.05). Spine types other than mushroom spines were unchanged among the groups (Table S3). These data suggest that CHI bilaterally decreased total hippocampal spine density on proximal and distal regions of CA1 apical dendrites. Spine loss was greater in the ipsilateral hippocampus. Additionally, CHI selectively lowered mushroom spine density. MN72 treatment limited the loss of mushroom spines as well as overall spine density.

### MN72 bilaterally maintains CA1 synaptic density after CHI

Preservation of spine density is suggestive of synapse preservation, yet Golgi-Cox visualizes only post-synaptic structures. Therefore, synapse density was measured by co-localization of the presynaptic marker synaptophysin with the postsynaptic marker PSD-95 (Figure 8A). Synaptophysin staining density had a significant group effect (F_4, 23_ = 7.12, p < 0.001) with CHI-Saline mice having less staining in the ipsilateral hippocampus than Sham-CHI and CHI-MN72 mice (p < 0.005; Figure 8B1). In contrast, synaptophysin staining density did not differ in either hippocampus in CHI-MN72 and Sham-CHI mice (p > 0.9). PSD-95 staining density did not significantly differ among groups (F_4, 23_ = 2.03, p > 0.1; Figure 8B1). Synaptophysin and PSD-95 immunostaining revealed a punctate distribution consistent with colocalization of presynaptic boutons and the PSD. Colocalization of synaptophysin and PSD-95 had a significant group effect (F_4, 23_ = 4.71, p < 0.01, Figure 8B2). Bilaterally, CHI-Saline mice had significantly fewer colocalized puncta as compared to Sham-CHI (ipsilateral, p < 0.05; contralateral, p < 0.005). In contrast, CHI-MN72 and Sham-CHI mice had similar amounts of colocalized puncta (p > 0.1). These data suggest that CHI lowered ipsilateral presynaptic termini and bilaterally reduced synaptic density. MN72 prevented the loss of presynaptic termini and the reduction of synaptic density.

**Figure 8.**
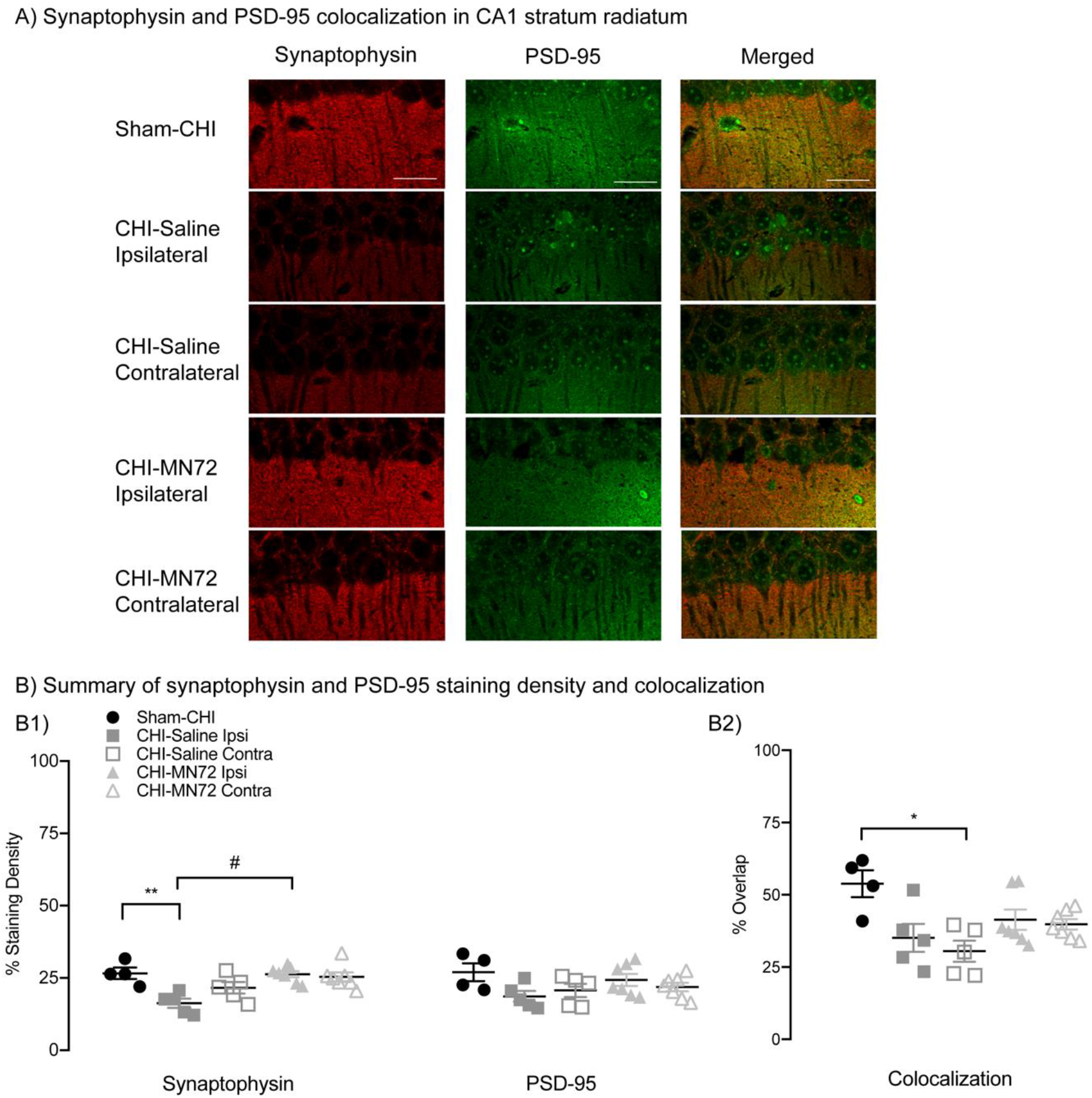
MN72 bilaterally maintains CA1 synaptic density after CHI. **A**, CA1 stratum radiatum stained for synaptophysin (red) and PSD-95 (green) from Sham-CHI, CHI-Saline and CHI-MN72 mice. **B**, Synaptophysin and PSD-95 density and colocalization. **B1**, Synaptophysin immunoreactivity in ipsilateral CA1 of CHI-Saline mice is lower than Sham-CHI and CHI-MN72 mice. PSD-95 immunoreactivity does not differ among the groups. **B2**, Synaptophysin and PSD-95 colocalization in CHI-Saline mice is decreased bilaterally compared to Sham-CHI mice; colocalization in CHI-MN72 and Sham-CHI mice does not differ. Scale bar, 50 μm, # represents significant difference with CHI-MN72, *p < 0.05, **p < 0.01, significant difference with Sham-CHI.

## Discussion

TBI drug development has largely focused on acute neuroprotective interventions that prevent or reduce the early consequences of TBI (Atkins, 2011). A limitation of this approach is few drugs have clinically useful therapeutic time windows. Drug efficacy rapidly diminishes due to the speed and evolving complexity of TBI pathophysiology, therefore we need a better understanding of what aspects of TBI can be improved by delayed drug administration (Mohamadpour et al., 2019). We took advantage of the long therapeutic time window of the drug combination MINO plus NAC to identify elements of TBI pathophysiology that could be targeted with drugs first dosed 72 hours after injury. CHI impairs performance on Barnes maze, which can be acquired with only one functional hippocampus (Klur et al., 2009; Shinohara et al., 2012). A first dose of MINO plus NAC at 24 hours PI did not prevent damage to the ipsilateral hippocampus (Sangobowale et al., 2018). MN72 treatment improved acquisition and retention of Barnes maze (Figure 1). Therefore, we examined both the ipsilateral and contralateral hippocampi of CHI-injured mice treated with saline or MN72 to determine whether MN72 targeted the contralateral hippocampus.

Unilateral CHI produced extensive bilateral damage both proximal and distal to the impact site with greater neuronal loss in the ipsilateral than in the contralateral hippocampus (Figure 2). Unilateral hippocampal damage is not sufficient to induce spatial memory impairment; rather bilateral damage is needed to produce deficits (Grady et al., 2003). Thus, it is likely a combination of neuroprotective effects in both hemispheres that underlies the functional improvements produced by MN72 treatment. Despite CA1 being more proximal to the impact site, CA3 neurons were more vulnerable to mechanical injury due to greater tensile strains that extend over a larger area than in CA1 (Casella et al., 2014). CA1 neurons, in contrast, sustained dendritic damage without cell death. Consistent with these observations, CHI bilaterally reduced CA3 neuronal density, while neuronal loss was observed only in ipsilateral CA1 (Figure 2). CA3 cell loss deafferents CA1, leading to CA1 dendritic arbor damage (Casella et al., 2014). Loss of CA3 afferents may contribute to disrupted dendritic architecture and reduced spine density in CA1. Selective neuronal loss demonstrates that hippocampal subregions support specific memory functions. CA3 lesions suggest that CA3 maintains short-term spatial representations, which are critical to spatial memory formation and recall (Gilbert and Brushfield, 2009; Le Duigou et al., 2014; Cherubini and Miles, 2015). Selective CA1 neuronal loss correlates with spatial learning deficits after injury (Auer et al., 1989; Milani et al., 1998). Injury to both CA3 and CA1 likely contributes to the learning and memory deficits produced by CHI (Figure 1).

Two complementary analyses characterized neuronal morphology and dendritic architecture on surviving neurons (Table S2). Bilateral damage to dendrites, spines and synapses was more extensive than neuronal loss, suggesting neuronal loss underestimates hippocampal injury. Surviving ipsilateral neurons were smaller and sustained more dendritic damage than corresponding contralateral neurons (Figures 3–5). In contralateral CA1, dendritic degeneration and synapse loss occurred despite minimal cell death (Figures 5, 8). CHI bilaterally lowered CA1 spine and synaptic density (Figures 7,8; Table S3). Reduced synaptophysin expression suggested additional selective loss of ipsilateral presynaptic termini. Thus, assessment of drug efficacy by neuroprotection alone is insufficient due to widespread damage to surviving neurons. The broader characterization of brain damage following CHI in this study uncovered elements of TBI pathophysiology that can be targeted by drugs first given days after injury.

MN72 prevented neuronal loss, protected neuronal morphology and increased spine and synapse density (Figures 2–8). MN72 bilaterally maintained CA1 dendritic arborization and complexity, suggesting that CA1 dendritic architecture can be treated with delayed drug administration. In contrast, in CA3 MN72 therapeutic effects were limited to the contralateral hippocampus. Ipsilateral CA3 may have been more severely damaged, or the kinetics of pathophysiological events in CA3 differ or occur more rapidly than in CA1. Recent experiments suggest that MN12 bilaterally maintains CA3 neuronal density (Whitney and Bergold, unpublished result). The differing effects of MN12 and MN72 dosing suggests more rapid pathological processes in ipsilateral than contralateral CA3.

Loss of the dendritic cytoskeletal protein MAP2 likely contributed to dendritic arbor damage. MAP2 degradation occurs rapidly following moderate TBI, and contributes to behavioral deficits (Taft et al., 1992). Previous studies demonstrated MAP2 loss in ipsilateral CA3 persisted for 1-month post-CHI (Grin’kina et al., 2016; Sangobowale et al., 2018). The additional analyses of hippocampal MAP2 expression in this study revealed regional differences in MAP2 loss and recovery. MAP2 loss occurred more rapidly in the ipsilateral than the contralateral hippocampus (Figure 6). In CA3, MAP2 loss was more rapid in basal dendrites, while in CA1 bilateral MAP2 loss occurred in both apical and basal dendrites within 3 days post-CHI (Figures 6). Similarly, MN72 therapeutic effects on MAP2 recovery differed by region. MN72 prevented MAP2 loss in contralateral CA3 basal dendrites (Figure 6B2), and accelerated or restored MAP2 levels in contralateral CA3 apical (Figure 6C2), CA1 basal (Figure 6D2), and CA1 apical dendrites (Figure 6E2). Decreased MAP2 degradation may result in MAP2 recovery since the injured neuron can reorganize and repair cytoskeletal structure (Taft et al., 1992). These results demonstrate distinct regional differences in the therapeutic window of MN72 for MAP2 expression. A recent study showed that L-cysteine prevented MAP2 loss in a weight drop TBI model, therefore NAC may underlie MN72 therapeutic effects on MAP2 expression (Ouyang et al., 2019).

In CA1 apical dendrites, Schaffer collaterals synapse on proximal branches, while perforant path axons make synapses on distal branches (Spruston, 2008). CHI bilaterally reduces dendritic complexity and spine density in both proximal and distal branches of apical dendrites (Figure 4, 5, 7). This likely disrupts dendritic integration of Schaffer collateral and perforant path inputs. MN72 bilaterally increased mushroom spine density on proximal branches. On distal dendrites, mushroom spine density increased only in the contralateral hippocampus. Mushroom spines are more stable than other spine types (Rodriguez et al., 2008). During memory formation, mushroom spine head size increases due to the transport of intracellular glutamate receptors that are essential for synaptic plasticity (Sebastian et al., 2013; Avila et al., 2017). Based on these properties, mushroom spines are hypothesized to be a physical substrate of long-term memories (Bourne and Harris, 2007). Restoration of spatial navigation and memory may arise, in part, by preserving dendritic geometry and mushroom spine density in the contralateral hippocampus. These data show that in addition to the structural integrity of the dendritic arbor, spine and synapse density can be successfully targeted with drugs first dosed days after injury.

The data in this study suggest that effective therapeutic interventions will target multiple injury mechanisms to reduce progressive secondary injury. Inflammation, mitochondrial damage and oxidative stress are secondary injuries that contribute to TBI pathogenesis and neuronal death (Blennow et al., 2012). Targeting multiple mechanisms likely results in a longer therapeutic window then targeting a single acute neuroprotective mechanism (Margulies et al., 2009). MINO plus NAC likely has multiple drug targets that underlie its many therapeutic actions. MINO plus NAC modulates inflammation (Chen et al., 2008; Garrido-Mesa et al., 2013; Haber et al., 2013; Eakin et al., 2014). MINO plus NAC may also reduce mitochondrial damage and oxidative stress. NAC restored mitochondrial dysfunction and brain glutathione levels after experimental TBI (Xiong et al., 1999). Neuroprotection on mitochondrial function may be related to the ability of NAC to directly scavenge free radicals and maintain mitochondrial calcium homeostasis (Eakin et al., 2014).

The CHI model used in this study reproduces many features of clinical TBI, however no TBI models recapitulate the complexity of human brain injury (Xiong et al., 2013). A limitation of most TBI models is the use of anesthesia during the injury procedure. Isoflurane, the anesthetic drug used in this study, may be neuroprotective, which could influence injury severity, pathology and recovery (Statler et al., 2006). This study only used male mice and therefore does not address sex as a biological variable. Sex differences in recovery outcomes and response to pharmacological therapies after injury are demonstrated both clinically and in TBI models (Wright et al., 2014; Failla and Wagner, 2015). In addition, this study assesses the effects of the drug combination MINO plus NAC, but not the individual drugs. MINO or NAC were previously shown to have little potency when dosed as individual drugs at 24 hours post-injury (Sangobowale et al., 2018). Ongoing work in our laboratory is investigating these issues in TBI pathophysiology, outcomes and therapeutic efficacy.

A major finding of this study is that MN72 has a different therapeutic time window in discrete hippocampal regions. Both drugs readily enter the brain making it unlikely that pharmacokinetics is responsible for regional therapeutic effects (Diaz-Arrastia et al., 2014). It is more likely that the complexity of TBI pathophysiology underlies the regional effects of MN72. Discrete brain regions may express the same TBI pathophysiology with different kinetics. Alternatively, discrete brain regions may vary in TBI pathophysiology. Either of these possibilities help explain why the therapeutic time window differs among different brain regions. Most preclinical studies have focused on drug therapeutic action proximal to the injury site, which resulted in the development of drugs with short therapeutic windows (Mohamadpour et al., 2019). Increased understanding of TBI pathophysiology suggests that a therapeutic time window is likely not limited to the first few hours after injury. TBI progresses over time, therefore delayed treatment may benefit TBI patients who cannot be treated immediately after injury.

A key unanswered question is what constitutes a clinically relevant therapeutic window for TBI intervention. Enrollment data for recent Phase III clinical trials show that TBI patients at designated trauma centers can be treated within 4-7 hours after a moderate to severe TBI (Skolnick et al., 2014). The 4-7 hour therapeutic window is optimal for patients who are injured enough to seek immediate medical attention. Less specialized hospitals will take longer to initiate treatment, and patients with mild TBI often wait longer to seek treatment after injury (Alexander, 1995; Kushner, 1998). To treat the largest number of patients, therapeutic interventions will need to retain high efficacy with delayed dosing. The finding that distal injury can be targeted with drugs first given days after injury provides a new framework in developing and evaluating drugs with sufficiently favorable therapeutic windows to treat TBI. The findings of this study provide further impetus to study the protective and restorative mechanisms of MINO plus NAC.

## Abbreviations

CHI: closed head injury
Contra: contralateral
Ipsi: ipsilateral
MAP2: microtubule associated protein 2
MINO: minocycline
MN12: MINO plus NAC first dosed 12 hours after injury
MN72: MINO plus NAC first dosed 72 hours after injury
NAC: N-acetylcysteine
TBI: Traumatic brain injury

## Supplemental Tables

**Table S1.** Barnes maze performance parameters. On training days 1-4, latency, errors, path length and speed are the average of trials 1-4 for each day. For the probe trial on day 5, percentage of time spent or holes searched in the target quadrant assessed recall of the escape box location.

| Acquisition Trials |  |  |  |  |
| --- | --- | --- | --- | --- |
| Day | Parameter | Sham-CHI | CHI-Saline | CHI-MN72 |
| 1 | Latency | 120.7±18.2 | 126.3 ± 2.2 | 121.4 ± 8.0 |
|  | Errors | 9.6 ± 1.3 | 10.84 ± 1.0 | 9.2 ± 1.7 |
|  | Path Length | 2.7 ± 0.3 | 3.3 ± 0.4 | 3.6 ± 0.4 |
|  | Speed | 0.03 ± 0.004 | 0.03 ± 0.005 | 0.03 ± 0.005 |
| 2 | Latency | 68.9 ± 11.2 | 130.5 ± 5.7 | 83.1 ± 9.2 |
|  | Errors | 5.5 ± 0.9 | 10.0 ± 1.2 | 5.5 ± 1.0 |
|  | Path Length | 2.4 ± 0.2 | 3.8 ± 0.6 | 2.6 ± 0.3 |
|  | Speed | 0.05 ± 0.008 | 0.04 ± 0.005 | 0.04 ± 0.005 |
| 3 | Latency | 37.2 ± 9.1 | 96.7 ± 10.3 | 74.7 ± 10.1 |
|  | Errors | 4.1 ± 1.2 | 7.0 ± 1.0 | 5.7 ± 1.0 |
|  | Path Length | 1.4 ± 0.5 | 2.8 ± 0.6 | 3.1 ± 0.6 |
|  | Speed | 0.05 ± 0.017 | 0.04 ± 0.003 | 0.07 ± 0.019 |
| 4 | Latency | 21.3 ± 2.8 | 85.1 ± 9.3 | 41.8 ± 3.7 |
|  | Errors | 2.1 ± 0.4 | 5.9 ± 0.4 | 3.0 ± 0.6 |
|  | Path Length | 1.3 ± 0.3 | 2.6 ± 0.6 | 1.5 ± 0.5 |
|  | Speed | 0.11 ± 0.035 | 0.05 ± 0.008 | 0.05 ± 0.017 |
| Probe Trial |  |  |  |  |
| 5 | % Time | 66.5 ± 5.3 | 36.1 ± 9.7 | 81.5 ± 6.6 |
|  | % Holes | 73.3 ± 6.7 | 22.5 ± 4.5 | 65.7 ± 11.3 |
|  | Distance | 3.4 ± 0.6 | 2.8 ± 0.5 | 2.3 ± 0.6 |

**Table S2.**
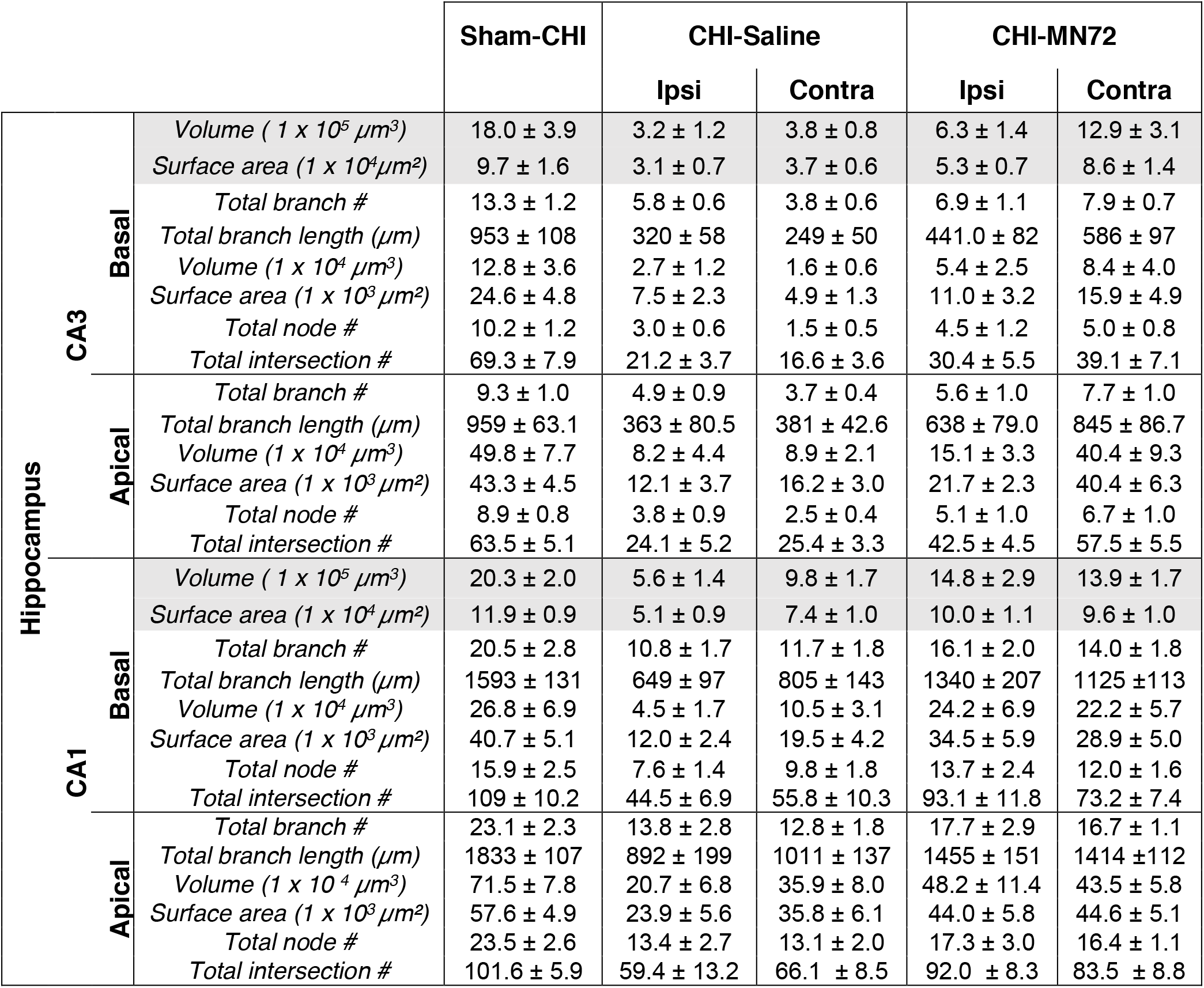
Morphological parameters of CA3 and CA1 pyramidal neurons are presented by treatment and hemisphere. All values are mean ± SEM.

**Table S3.**
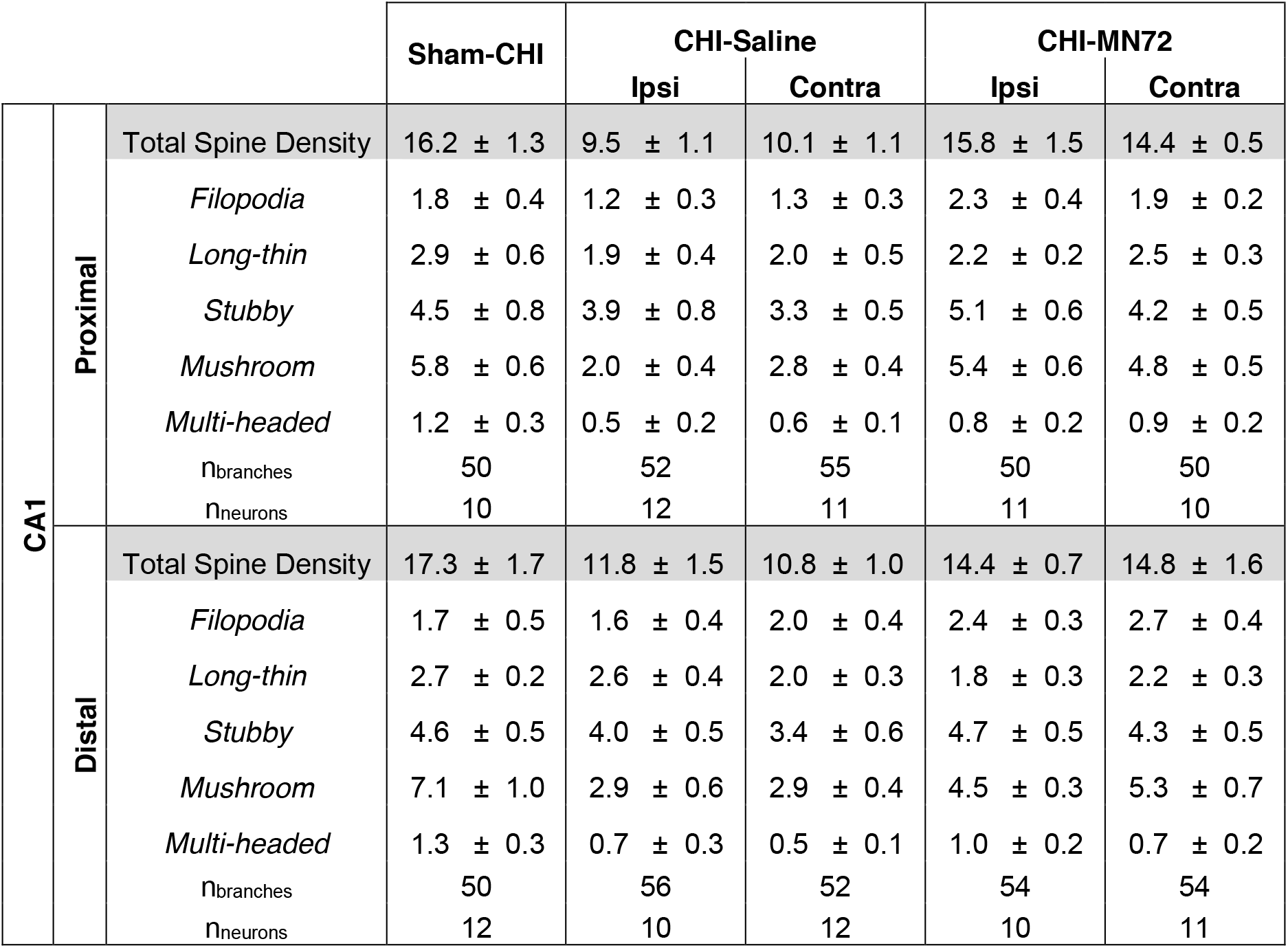
Analysis of proximal and distal CA1 apical dendrites for total spine density and spine density by morphological subtype. All values are mean ± SEM.

| CA1 |  |  | Sham-CHI | CHI-Saline |  | CHI-MN72 |  |
| --- | --- | --- | --- | --- | --- | --- | --- |
|  |  |  |  | Ipsi | Contra | Ipsi | Contra |
| CA1 | Proximal | Total Spine Density | 16.2 ± 1.3 | 9.5 ± 1.1 | 10.1 ± 1.1 | 15.8 ± 1.5 | 14.4 ± 0.5 |
|  |  | *Filopodia* | 1.8 ± 0.4 | 1.2 ± 0.3 | 1.3 ± 0.3 | 2.3 ± 0.4 | 1.9 ± 0.2 |
|  |  | *Long-thin* | 2.9 ± 0.6 | 1.9 ± 0.4 | 2.0 ± 0.5 | 2.2 ± 0.2 | 2.5 ± 0.3 |
|  |  | *Stubby* | 4.5 ± 0.8 | 3.9 ± 0.8 | 3.3 ± 0.5 | 5.1 ± 0.6 | 4.2 ± 0.5 |
|  |  | *Mushroom* | 5.8 ± 0.6 | 2.0 ± 0.4 | 2.8 ± 0.4 | 5.4 ± 0.6 | 4.8 ± 0.5 |
|  |  | *Multi-headed* | 1.2 ± 0.3 | 0.5 ± 0.2 | 0.6 ± 0.1 | 0.8 ± 0.2 | 0.9 ± 0.2 |
|  |  | n<sub>branches</sub> | 50 | 52 | 55 | 50 | 50 |
|  |  | n<sub>neurons</sub> | 10 | 12 | 11 | 11 | 10 |
|  | Distal | Total Spine Density | 17.3 ± 1.7 | 11.8 ± 1.5 | 10.8 ± 1.0 | 14.4 ± 0.7 | 14.8 ± 1.6 |
|  |  | *Filopodia* | 1.7 ± 0.5 | 1.6 ± 0.4 | 2.0 ± 0.4 | 2.4 ± 0.3 | 2.7 ± 0.4 |
|  |  | *Long-thin* | 2.7 ± 0.2 | 2.6 ± 0.4 | 2.0 ± 0.3 | 1.8 ± 0.3 | 2.2 ± 0.3 |
|  |  | *Stubby* | 4.6 ± 0.5 | 4.0 ± 0.5 | 3.4 ± 0.6 | 4.7 ± 0.5 | 4.3 ± 0.5 |
|  |  | *Mushroom* | 7.1 ± 1.0 | 2.9 ± 0.6 | 2.9 ± 0.4 | 4.5 ± 0.3 | 5.3 ± 0.7 |
|  |  | *Multi-headed* | 1.3 ± 0.3 | 0.7 ± 0.3 | 0.5 ± 0.1 | 1.0 ± 0.2 | 0.7 ± 0.2 |
|  |  | n<sub>branches</sub> | 50 | 56 | 52 | 54 | 54 |
|  |  | n<sub>neurons</sub> | 12 | 10 | 12 | 10 | 11 |

## Acknowledgements

The authors thank Drs. Sheryl Smith, Julie Parato and Mr. Matthew Evrard for their assistance with the Golgi-Cox studies. This work was supported by NIH grant NS108190-01 (to P.J.B.)

